# The angiotensin antagonist Losartan modulates social reward motivation and punishment sensitivity via modulating midbrain-striato-frontal circuits

**DOI:** 10.1101/2021.07.19.452920

**Authors:** Xinqi Zhou, Ting Xu, Yixu Zeng, Ran Zhang, Ziyu Qi, Weihua Zhao, Keith M Kendrick, Benjamin Becker

## Abstract

Social deficits and dysregulations in dopaminergic midbrain-striato-frontal circuits represent transdiagnostic symptoms across psychiatric disorders. Animal models suggest that interactions between the dopamine and renin-angiotensin system may modulate learning and reward-related processes. The present study therefore examined the behavioral and neural effects of the angiotensin II type 1 receptor (AT1R) antagonist Losartan on social reward and punishment processing in humans. A pre-registered randomized double-blind placebo-controlled between-subject pharmacological design was combined with a social incentive delay fMRI paradigm during which subjects could avoid social punishment or gain social reward. Healthy volunteers received a single-dose of Losartan (50mg, n=43) or placebo (n=44). Reaction times and emotional ratings served as behavioral outcomes, on the neural level activation and connectivity were modelled. Relative to placebo, Losartan modulated the reaction time and arousal differences between social punishment and social reward. On the neural level the Losartan-enhanced motivational salience of social rewards was accompanied by stronger ventral striatum-prefrontal connectivity during reward anticipation. Losartan increased the reward-neutral difference in the ventral tegmental area (VTA) and attenuated VTA associated connectivity with the bilateral insula in response to punishment during the outcome phase. Losartan modulated approach-avoidance motivation and emotional salience during social punishment versus social reward via modulating distinct core nodes of the midbrain-striato-frontal circuits. The findings document a modulatory role of the renin-angiotensin system in these circuits and associated social processes, suggesting a promising treatment target to alleviate social dysregulations.

**Significance Statement:** Social deficits and anhedonia characterize several mental disoders and have been linked to the midbrain-striato-frontal circuits of the brain. Based on initial findings from animal models we here combine the pharmacological blockade of the angiotensin II type 1 receptor (AT1R) via Losartan with functional MRI to demonstrate that AT1R blockade enhances the motivational salience of social rewards and attenuates the negative impact of social punishment via modulating the communication in the midbrain-striato-frontal circuits in humans. The findings demonstrate for the first time an important role of the AT1R in social reward processing in humans and render the AT1R as promising novel treatment target for social and motivational deficits in mental disoders.

## Introduction

Adaptive processing of social feedback is vital for interpersonal functioning and mental health. Based on the Research Domain Criteria framework, dysregulations in these domains (social communication and reward/loss evaluation) and their underlying neural processes contribute to the development and maintenance of psychiatric disorders (Cuthbert and Insel, 2013). These domains may represent a transdiagnostic treatment target with the potential to improve social functioning. Dysregulations in midbrain-striato-prefrontal circuits have been increasingly established as a core pathogenic mechanism across psychiatric disorders (Der-Avakian and Markou, 2012; Russo and Nestler, 2013; Luijten et al., 2017; Fenster et al., 2018). Both human imaging and animal studies suggest that this circuitry involves in social reward and punishment processing (Dolen et al., 2013; Hung et al., 2017; Gu et al., 2019; Martins et al., 2021). Dopamine (DA) and its interactions with other neurotransmitter systems such as oxytocin play an important role in modulating social reward and punishment in these cicruits (Nawijn et al., 2017; Grimm et al., 2021a), however, direct pharmacological modulation of these systems commonly results in negative side effects or highly context-dependent effects, respectively, which critically impede the clinical utility of these approaches (Pessiglione et al., 2006; Grimm et al., 2021b; Quintana et al., 2021).

Recent pharmacological studies in healthy humans have demonstrated that targeting the renin-angiotensin system (RAS) via the angiotensin II type 1 receptor (AT1R) antagonist Losartan (an approved treatment for hypertension) can modulate reward and threat processing as well as learning and memory in the absence of negative side effects (Marvar et al., 2014; Reinecke et al., 2018; Pulcu et al., 2019; Zhou et al., 2019a; Swiercz et al., 2020). Earlier animal models suggest an interaction between the RAS and the central DA system, including a dense expression of RAS receptors in midbrain-striato-prefrontal circuits (Chai et al., 2000) and functionally significant angiotensin II receptors located presynaptically on dopaminergic neurons (Medelsohn et al., 1993; Brown et al., 1996). Losartan induced concentration-dependent inhibition of dopamine release via inactivation of AT1R (Narayanaswami et al., 2013), but also enhanced dopamine D1 receptor signaling which may contribute to both its effects on hypertension (Li et al., 2012) and reward-related processes (Maul et al., 2005; Hosseini et al., 2009). Together, the available evidence suggests that targeting the RAS via Losartan may represent a promising candidate to modulate neural processing in midbrain-striatal-prefrontal circuits which mediate earlier stages of social and non-social reward processing to improve behavioral adaption (Izuma et al., 2008; Wake and Izuma, 2017; Gu et al., 2019; Grimm et al., 2021a; Martins et al., 2021). Initial evidence for this strategy in humans comes from a recent study that demonstrated that a single dose of 50mg Losartan can modulate feedback-dependent learning in healthy individuals such that Losartan enhanced the difference between loss and reward feedback learning rates and suppressed loss learning rates (Pulcu et al., 2019). These findings align with previous studies reporting valence-specific effects of DA and may suggest a pathway via which losartan may amplify the balance between reward and punishment (Pessiglione et al., 2006; Esser et al., 2021).

Against this background we combined a pre-registered randomized double-blind between-group placebo-controlled pharmacological experiment with functional MRI (fMRI) to examine whether social reward and punishment processing can be modulated by a single dose of Losartan, thus bridging the translational gap between animal model and human research as well as to determine the clinical potential of Losartan. To this end healthy volunteers underwent a well-validated social incentive delay fMRI paradigm (Nawijn et al., 2017). Behavioral indices reflecting motivation and subsequent emotional impact of social feedback, neural indices during reward and punishment anticipation and outcomeserved as primary outcomes. Based on findings from animal and human studies we hypothesized that Losartan would (a) enhance differential processing of reward and punishment on the behavioral level (Pulcu et al., 2019), which on the neural level would be reflected in (b) enhanced differential activiation and connectivity in VTA-striatal-frontal circuits during social reward-punishment processing.

## Materials and method

### Participants

Ninety healthy participants (age range 18-27 years) were recruited for the randomized placebo-controlled between-subject pharmacological fMRI study which encompassed a single-dose p.o. administration of 50mg Losartan or placebo and subsequent administration of a social incentive delay fMRI paradigm (SID) with a demonstrated sensitivity to capture pharmacological modulations (Nawijn et al., 2017). N = 87 subjects (N=43, 26 males, Losartan; N=44, 24 males, placebo; **Table 1**) were included in the final analyses (exclusion see **supplementary method**). Given the complexity of the modelling and fMRI analyses the sample size was based on recent between-subject pharmacological studies employing samples of 35-40 subjects to determine behavioral and neural effects of Losartan (Zhou et al., 2019a; Shkreli et al., 2020).

**Table 1.**
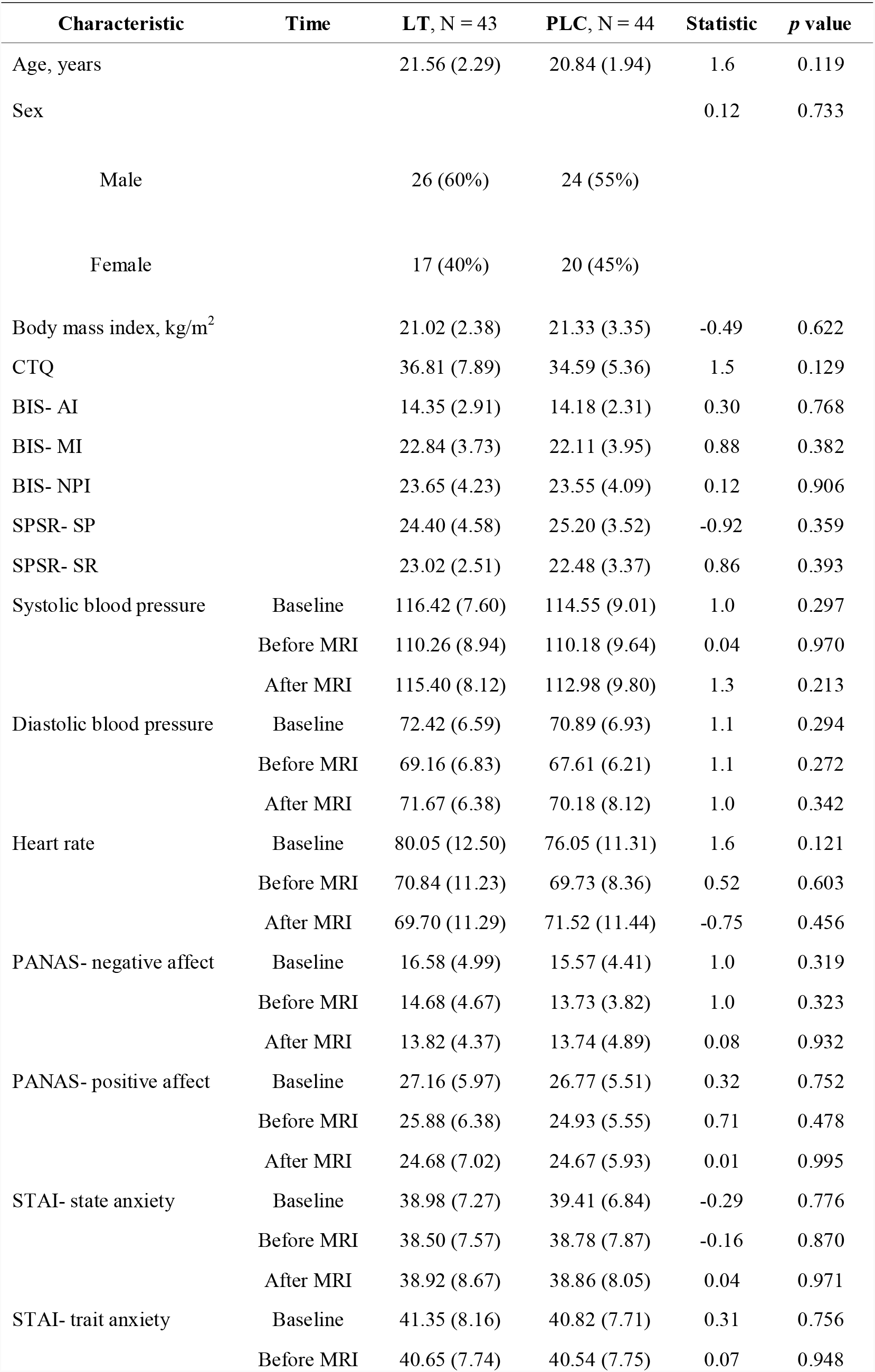
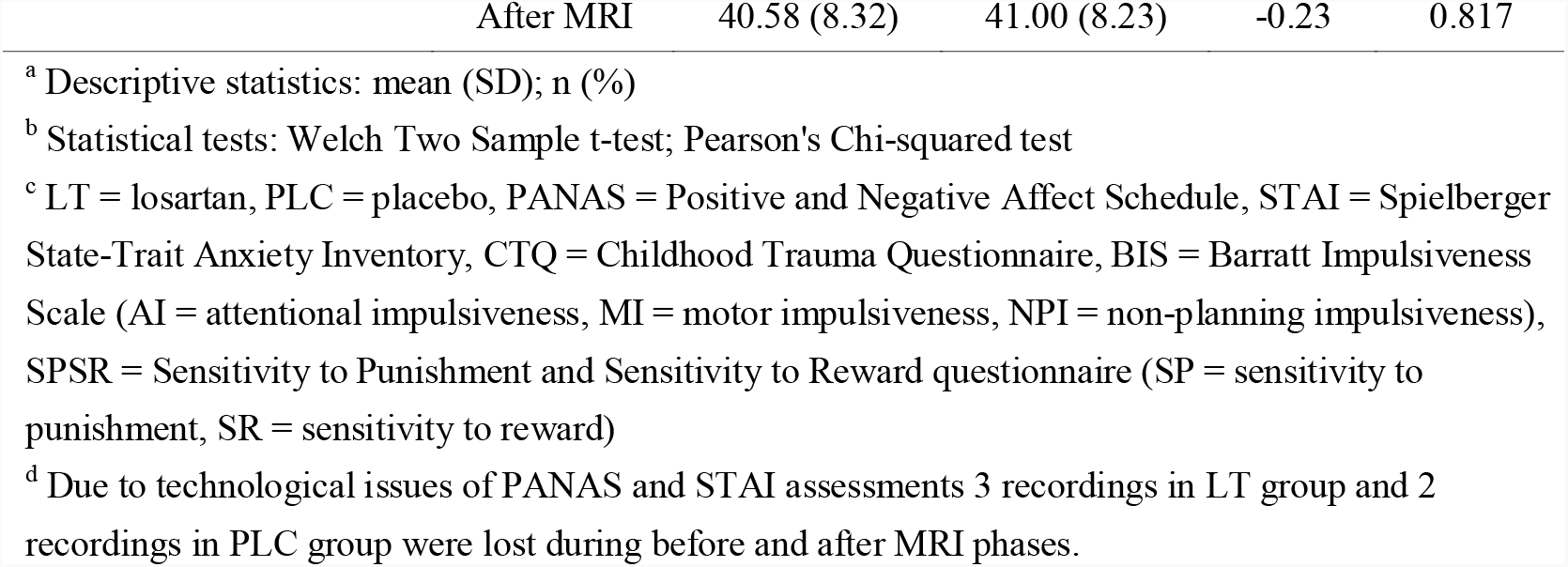
Participant Demographics and Control Measures

### Pharmacological and experimental procedure

Participants were stratified for sex and randomly allocated to treatment. Treatment was packed in identical capsules, counterbalanced across sexes and dispensed by an independent researcher. To reduce potential confounding effects of early life stress (Birn et al., 2017), impulsiveness, sensitivity to punishment and reward on reward-related neural processing the Childhood Trauma Questionnaire (CTQ), Barratt Impulsiveness Scale (BIS), and Sensitivity to Punishment and Sensitivity to Reward Questionnaire (SPSRQ) were administered at baseline (**Figure 1A**) (Barratt, 1959; Torrubia et al., 2001; Bernstein et al., 2003). Given that after oral administration Losartan peak plasma levels are reached after 90 minutes with a terminal elimination half-life ranging from 1.5 to 2.5 hours (Ohtawa et al., 1993; Lo et al., 1995; Sica et al., 2005) the experimental paradigm started 90 minutes after treatment (see also(Mechaeil et al., 2011; Zhou et al., 2019a)). Losartan rapidly crosses the blood-brain barrier (Li et al., 1993; Culman et al., 1999) and while effects at central receptors have been observed after 30 minutes after i.e. administration effects on cardiovascular indices only become apparent after 3 hours (e.g. (Ohtawa et al., 1993; Pulcu et al., 2019; Zhou et al., 2019a)). To further control for potential confounding effects of Losartan on cardiovascular activity blood pressure and heart rate were assessed before drug administration, as well as before and after the SID fMRI paradigm (**Figure 1A**, details **supplementary methods**). To control for nonspecific affective effects of Losartan the affective state of participants was tracked troughout the experiment via the Spielberger State–Trait Anxiety Inventory (STAI) and the Positive and Negative Affective Scale (PANAS) which were administered before drug administration, at the time of peak plasma concentrations and after the experiment (Spielberger et al., 1983; Watson et al., 1988). The subsequent affective impact of Losartan-induced changes on social feedback processing was assessed via ratings of the cues before treatment, after fMRI, and following feedback stimuli after fMRI (**Figure 1A**). After the entire experiment participants were asked to guess the treatment they received.

**Figure 1.**
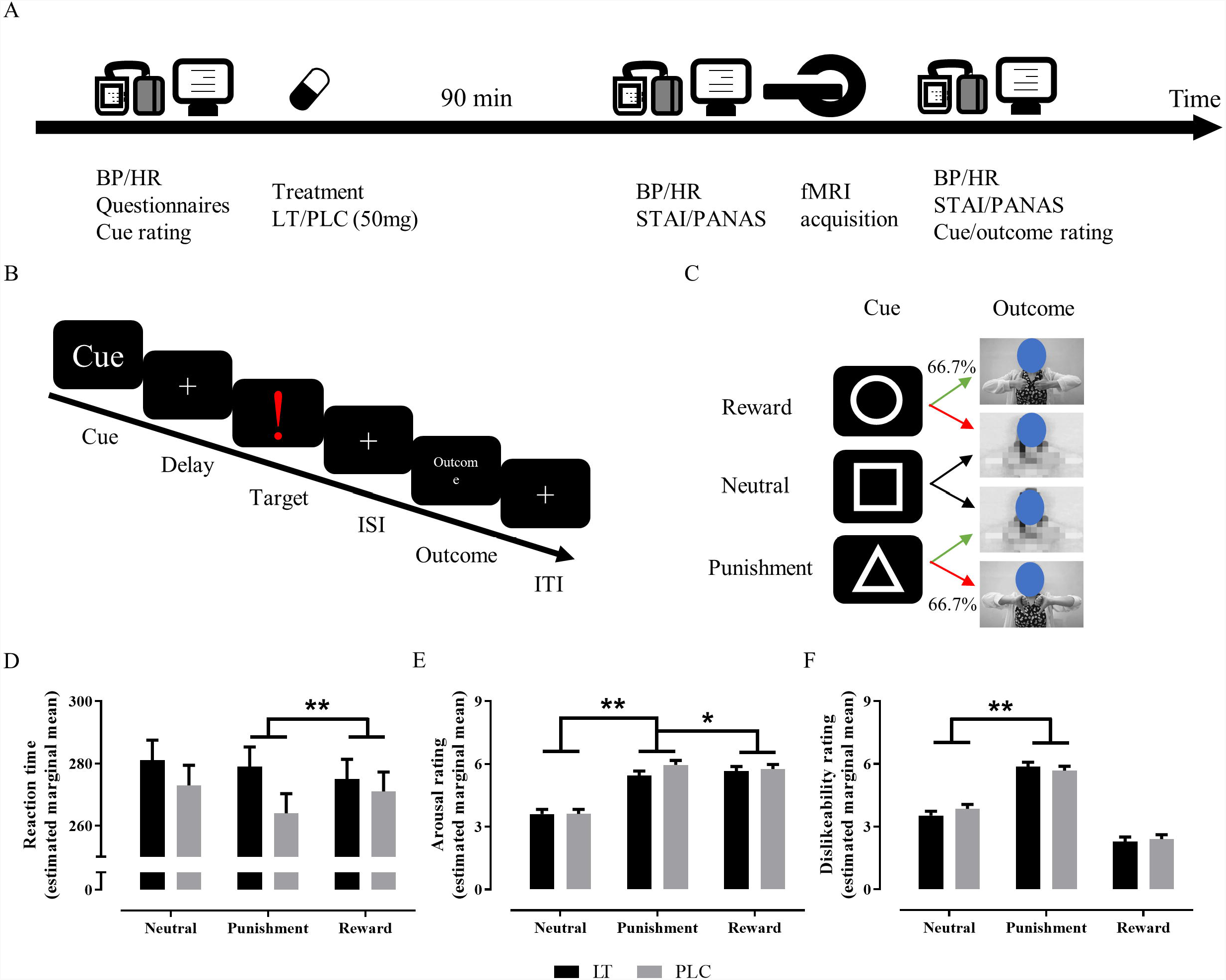
Schematic depiction of the experimental protocols and the experimental paradigm. (A) The entire experimental procedure encompassed baseline assessment, drug administration, assessments before fMRI (corresponding to estimated peak plasma levels) and post fMRI acquisition. Abbreviations: BP = blood pressure, HR = heart rate, LT = losartan, PLC = placebo, STAI = state-trait anxiety inventory, PANAS = positive and negative affect schedule. (B) Schematic representation of the trial structure in the SID paradigm. Each trial started with a 500ms cue presentation (circle, reward trial; square, neutral trials, and triangle, punishment trials) followed by a delay (jittered between 1000 and 3000 ms). Next the target appeared with a duration adjusted to the individual response time. After an inter-stimulus interval (ISI, 2000 ms – target duration) the outcome was presented for 1500 ms, followed by an inter-trial interval (ITI) with a duration jittered between 1000 and 3000 ms. (C) Cues and corresponding outcomes (for display only) per trial type. By means of an adaptive algorithm that increased or decreased target durations the rate of reward and punishment feedback was adapted across subjects. This ensured that 66.7% of reward-cue and punishment-cue trials were followed by social reward or punishment, respectively. The neutral cue was always followed by neutral feedback. Estimated marginal mean and stand error of reaction time (D), arousal rating (E), and dislikeability rating (F) are presented. * and ** denote relevant significant post hoc differences at *p* < 0.05 and *p* < 0.01 respectively.

Written informed consent was obtained, the study was approved by the local institutional ethics committee, all procedure followed the Declaration of Helsinki, and all procedures were preregistered (NCT04604756, URL: https://clinicaltrials.gov/ct2/show/NCT04604756). In the pre-registartion modulatory effects on the VTA-striato-frontal brain activity served as primary outcomes while effects on the behavioral level served as secondary outcomes. The analyses on the network level and the neural PE signal were not pre-registered and are thus exploratory in nature.

### Social incentive delay task

We employed a validated social incentive delay (SID) fMRI paradigm (Nawijn et al., 2017). The paradigm presents condition-specific cues (positive, negative, neutral) which signal that a social reward can be obtained or a social punishment can be avoided (anticipation). Next, participants undergo a reaction time task which is followed by the presentation of a possibility-dependent social reward, punishment or neutral feedback (outcome) (**Figure 1B-C**). To facilitate a balanced and sufficient number of trials to support a robust analysis on the neural level the number of trials for each outcome was adopted by means of an adaptive algorithm (details see supplement).

### Behavioral analysis

To maintain the trial-specific information of the SID task and increase sensitivity, a linear mixed model was used with condition (social reward, punishment, neutral) and treatment (Losartan, placebo) as two fixed factors and subject as random factor to account for individual adaptations of reaction time windows. Trials with no responses and RTs ±3SD on the individual level were removed. Treatment effects on emotional perception ratings of cues and outcomes were examined with separate ANOVA and liner mixed models (see **supplementary method**).

### MRI data acquisition and preprocessing

MRI data was acquired using a 3T GE MR750 system. Preprocessing was fully implemented in fMRIPrep (Esteban et al., 2019) except for smoothing with a Gaussian kernel at full width at half maximum (FWHM, 8×8×8mm) conducted in SPM12 (Welcome Department of Cognitive Neurology, London, UK, http://www.fil.ion.ac.uk/spm/software/spm12) (Friston et al., 1994). Details see **supplementary method**.

### Individual- and group-level BOLD fMRI analyses

On the first-level the SID task was modeled in line with previous studies (Rademacher et al., 2010; Lawn et al., 2020; Zhang et al., 2020; Faulkner et al., 2021), including condition-(social reward, social punishment, neutral) and phase-(anticipation, outcome) specific regressors (see **supplementary method**). On the group level effects of treatment were examined employing mixed ANOVA analyses with the factors (condition, treatment) for each phase. Based on our a priori regional hypotheses the analyses focused on the ventral striatum (VS), dorsal striatum (DS) (based on the brainnetome atlas, see also Zhou et al., 2019b) and the unthresholded probabilistic masks of the VTA from a VTA atlas (Trutti et al., 2021) as Regions of Interest (ROIs).

### Exploratory functional connectivity analysis

No effect of condition was observed during anticipation in our a priori defined network encompassing the VS/DS/VTA (Figure S1). To determine the social reward-punishment networks we next examined neural activity during receipt of feedback [reward+punishment–neutral] in the entire sample. Results revealed that social feedback induced stronger activity in regions involved in salience, value and social processes, including insula, striatum, dorsal medial prefrontal cortex (dmPFC), and occipital lobe (**Figure S1C**). Combined with the a priori defined VTA-striatal structural masks three peak coordinates (VS: [22/-6/-10], DS: [-14/-2/-8], VTA: [10/-14/-12]) were identified to construct spherical seeds with 6 mm radius which served as seeds for the generalized psychophysiological interactions (gPPI) analysis (McLaren et al., 2012) (**supplementary method**). Treatment effects were determined by comparing the seed-region-specific connectivity maps by means of mixed ANOVA analyses with the factors (condition, treatment) on the whole brain level (separately for each phase). To further disentangle significant interaction effects parameter estimates were extracted from regions exhibiting significant interaction effects involving treatment.

### Thresholding

ROI analyses were conducted in the R package ‘afex’, and the statistical significance level set to *p*<.05. On the whole-brain level an initial cluster-forming threshold was set to voxel level *p*<.001, and statistical significance was determined via cluster-level inference and familywise error (FWE) control for multiple comparisons with *p*_*FWE*_<.05 (Slotnick, 2017).

## Results

### Participants

Treatmemnt groups (losartan, n=43; placebo, n=44) exhibited comparable sociodemographic and psychometric characteristics (**Table 1**). During the experiment no differences in baseline assessments or changes in heart rate, blood pressure, and emotional state were observed between the treatment groups and total guess accuracy was 52.87% together arguing against the impact of potential confounders and unspecific effects of Losartan.

### Losartan effects on reaction time and affective response to outcome stumuli

The linear mixed model revealed a significant interaction effect (*F*=3.706, *p*=0.025, **Figure 1D**, Table S1) between condition and treatment on reaction times indicating that Losartan induced significantly stronger differences between social punishment vs social reward as compared to placebo (*t*=2.679, *p*=0.007), reflecting a possibility of Losartan mediated the approach-avoidance motivation of social feedback. The main effects of treatment and condition were not significant.

Examining effects of the experimental manipulation and treatment on the affective evaluation by means of a linear mixed model revealed a significant condition main (*F*=404.983, *p*<0.0001, **Figure 1E**) and condition times treatment interaction effect (*F*=4.914, *p*=0.007) on arousal ratings for outcomes. Post hoc analyses showed that Losartan increased the reward-punishment difference (*t*=2.390, *p*=0.017) and decreased punishment-neutral difference (*t*=-2.952, *p*=0.003) relative to placebo. With respect to dislikeability ratings for the outcomes a linear mixed model revealed significant condition main (*F*=633.848, *p*<0.0001, **Figure 1F**), and condition times treatment interaction effects (*F*=3.413, *p*=0.033). Post hoc tests showed that Losartan increased the punishment-neutral difference (*t*=2.597, *p*=0.0095) relative to placebo. No significant treatment main or interaction effects were observed on cue or other outcome ratings (see **supplementary results**).

### Losartan effects on neural activation during anticipation and outcome phase

No significant main or interaction effects of treatment were observed in the a priori ROI analyses on extracted parameter estimates during anticipation. An exploratory whole-brain analysis confirmed the lack of significant treatment main and interaction effects (at *p*_*FWE*_<0.05). During the outcome phase a significant treatment times condition effect was observed in the VTA (*F*=3.24, *p*=0.0435), reflecting that Losartan significantly increased the difference between reward and neutral (*t*=2.407, *p*=0.0172) as well as between reward and punishment (*t*=1.924, *p*=0.056, marginal significant, see **supplementary results**). Significant main effects of condition during the anticipation and outcome phase see **supplementary results** (**Figure S1, Table S2**).

### Losartan effects on VS-networks during anticipation

On the network level significant interaction effects between condition and treatment were found for the VS during anticipation but for the VTA during the outcome phase (all findings passed whole-brain *p*_*FWE*_<0.05, **Figure 2, Table 2**). Subsequent post-hoc tests revealed that Losartan significantly modulated VS-middle frontal gyrus (MFG) connectivity during neutral-punishment (*t*=2.541, *p*=0.0119), punishment-reward (*t*=-3.910, *p*=0.0001), and between social reward feedback (*t*=2.451, *p*=0.0151) processes, with the effects being driven by enhanced coupling during social reward-anticipation.

**Table 2.** Functional connectivity results

| Cluster region | Cluster size | x | y | z | *F* value |
| --- | --- | --- | --- | --- | --- |
| Anticipation phase, VS seed |  |  |  |  |  |
| MFG | 309 | 38 | 56 | 8 | 13.69 |
|  |  | 38 | 60 | 16 | 11.41 |
| Outcome phase, VTA seed |  |  |  |  |  |
| L insula, STG | 427 | -44 | -22 | -2 | 15.16 |
|  |  | -44 | -6 | 0 | 12.50 |
|  |  | -40 | 6 | -6 | 10.44 |
| R insula, putamen | 246 | 40 | 12 | 16 | 14.09 |
|  |  | 38 | 4 | -6 | 11.07 |
|  |  | 30 | 12 | 10 | 10.93 |
| SFG | 213 | -18 | 20 | 56 | 11.75 |
|  |  | -8 | 30 | 60 | 9.83 |
|  |  | -30 | 22 | 46 | 8.26 |

Note: All clusters passed the threshold at whole-brain cluster level $p_{FWE} < .05$ . L = left, R = right, SFG = superior frontal gyrus, MFG = middle frontal gyrus, STG = superior temporal gyrus.

**Figure 2.**
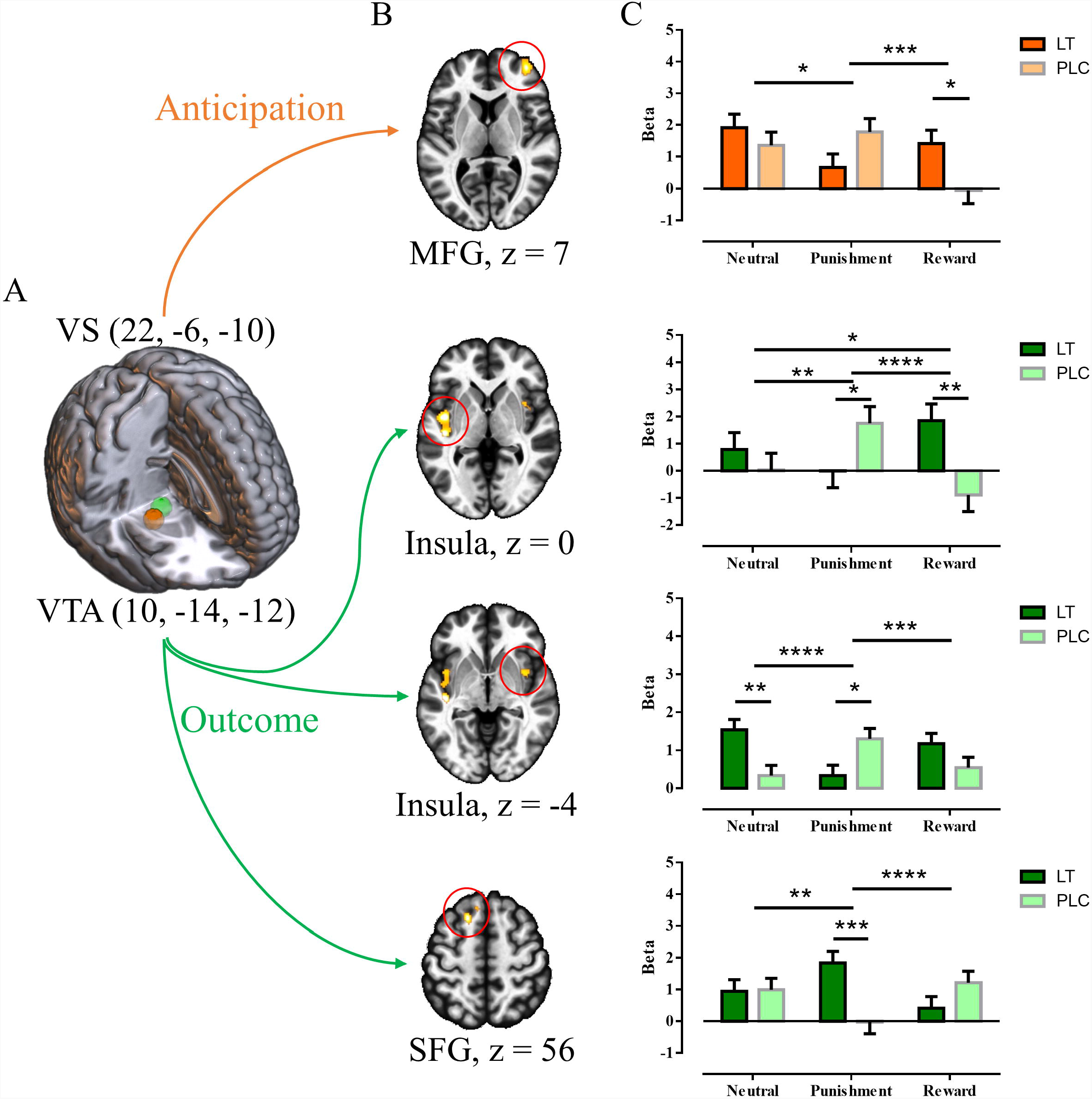
Effects of Losartan on the network level. (A) Seeds of interest, i.e. VS and VTA. (B) Regions exhibiting significant conditions times treatment interaction effects during anticipation and outcome phases. (C) Post hoc tests on extracted parameters from each significant clusters. VS = ventral striatum VTA = ventral tegmental area, SFG = superior frontal gyrus, MFG = middle frontal gyrus, LT = losartan, PLC = placebo, *, **, ***, and **** denote relevant significant post hoc differences at *p* < 0.05, *p* < 0.01, *p* < 0.001, and *p* < 0.0001 respectively.

### Losartan effects on VTA-networks during outcome

In contrast, Losartan specifically modulated VTA-networks during outcome, such that Losartan modulated VTA-insula (left) connectivity during the neutral-punishment pattern (*t* =2.613, *p*=0.0098), the punishment-reward pattern (*t*=-4.671, *p*<0.0001), the neutral-reward pattern (*t*=-2.059, *p*=0.0410), and within social punishment (*t*=-2.012, *p*=0.0456) and social reward (*t*=3.128, *p*=0.002) respectively. In addition, Losartan modulated both VTA-insula (right) and VTA-SFG connectivity for neutral-punishment(*t*=5.023, *p*<0.0001; *t*=-3.127, *p*=0.0021) and punishment-reward patterns (*t*=-3.683, *p*=0.0003; *t*=4.368, *p*<0.0001, **Figure 2C**). Losartan also changed VTA-insula (right) connectivity in social punishment (*t*=-2.512, *p*=0.0128) and neutral (*t*=3.13, *p*=0.002), VTA-superior frontal gyrus (SFG) connectivity in social punishment (*t*=3.613, *p*=0.0004). A direct comparion between treatments revealed consistent effects of Losartan on processing of social punishment feedback, such that it decreased VTA communication with the bilateral insula, yet enhanced VTA communication with the SFG.

## Discussion

The present pharmacological fMRI trial aimed to determine whether targeting the RAS system via Losartan can modulate social reward and punishment processing via modulating VTA-striatal-frontal circuits. On the behavioral level Losartan modulated the motivational significance of social reward and punishment during anticipation while affecting the subsequent affective evaluation of social stimuli. On the neural level the enhanced motivational significance was reflected by increased coupling between the VS and MFG during anticipation of social rewards. During the outcome phase Losartan enhanced neural signals of the reward-neutral difference in the VTA while attenuating VTA-insula communication and concomitantly enhancing VTA-SFG communication during social punishment. Notably, several of our pre-registered predictions with respect to a Losartan-induced modulation of regional brain activity during the anticipation and outcome of social reward and punishment were not confirmed (except effects on the VTA). Instead, the results from the exploratory network level analyses provided a more complex picture of the regulatory effects of Losartan on VTA-striato-frontal communication during reward-related processes. Together, these findings provide first insights into the regulatory role of the renin angiotensin system on meso-striato-cortical pathways that may underly effects in the domain of punishment and reward processing.

On the behavioral level Losartan modulated the motivational significance and arousal experience for social punishment relative to social reward feedback, an effect that was mainly driven by prolonged reaction times during anticipation of and subsequently reduced arousal reaction towards social punishment stimuli. These findings partly align with observations in previous studies, such that following Losartan healthy subjects perceived loss outcomes as being less informative resulting in an attenuated loss learning rate (Pulcu et al., 2019), and exhibited accelerated extinction and autonomous arousal decreases towards threat (Zhou et al., 2019a). Together, these observations indicate that Losartan may attenuate the impact of negative information thus promoting a relative higher influence of anticipatory motivation and post encounter learning towards positive information.

On the neural level the modulation of the approach-avoidance motivation between negative and positive social information was accompanied by a modulation of VS-frontal circuits, such that Losartan reduced VS-MFG connectivity during anticipation of social punishment but increased connectivity in this circuit during anticipation of social reward. Convergent evidence suggests that the VS plays a key role in dopamine-mediated anticipatory and motivational processes (Izuma et al., 2008; Gu et al., 2019; Martins et al., 2021) and that the pathways between the VS and frontal regions are critically involved in associated social processes including motivational and reinforcing aspects of social interactions (Murugan et al., 2017; Modi and Sahin, 2019). In patients with marked social impairments pharmacological modulation of the coupling between VS and MFG has been associated with improved computation of future positive social outcomes (Gordon et al., 2016; Greene et al., 2018) and effects on this circuit may thus reflect a potential mechanism via which Losartan can increase social motivation.

In contrast to the modulation of VS-centered circuits during the anticipation stage, Losartan specifically modulated VTA activity as well as its connectivity with insular and frontal regions during the outcome phase. During the social feedback presentation stage Losartan increased the differential processing of rewarding feedback from both, negative as well as neutral feedback in the VTA. The VTA represents a pivotal node in dopamine-modulated reward processing and learning circuits (Averbeck and Costa, 2017; Sharpe et al., 2017; Gu et al., 2019; Grimm et al., 2021a) and together with the amygdala drives dopaminergic signaling in response to social stimuli (Modi and Sahin, 2019; Grimm et al., 2021a), suggesting that Losartan rendered positive social signals as more salient. In contrast, Losartan specifically decreased coupling of the VTA with the bilateral mid-posterior insula in response to social punishment. The insula plays a key role in salience and interoceptive information processing, with the mid-posterior insula being involved in representing the intensity of aversive experiences (Uddin, 2015; Zhou et al., 2020). This suggests that Losartan may have attenuated the aversive emotional impact of negative social feedback on the insula leading to lower arousal ratings for the negative social stimuli following the experiment.

From a functional neuroanatomy perspective Losartan modulated neural activity and connectivity of distinct key nodes of the midbrain-striatal system during different aspects of social feedback processing. Thus, VS connectivity was specifically affected during anticipation while VTA networks. This dissociation aligns with the distinct functions of these core nodes in feedback-associated social and non-social processes (Zhou et al., 2019b; Gordon et al., 2021; Suzuki et al., 2021). The VTA encompasses the majority of dopaminergic cell bodies and is strongly involved in predicting outcomes including social error signals and guiding flexible adaptation (Birn et al., 2017; Hetu et al., 2017; Gu et al., 2019; Grimm et al., 2021a), whereas the VS which receives dopaminergic projections from the VTA, is strongly involved in appetitive motivation and reward-expectation for both social and non-social feedback (Gu et al., 2019; Martins et al., 2021) while the DS is stronger involved in learning, action initiation, and habit formation (Klugah-Brown et al., 2020; Suzuki et al., 2021).

Altough most of these functions encompass social as well as non-social processes their critical role in reward and punishment processing critically influences social behavior (Murugan et al., 2017; Modi and Sahin, 2019; Zhang and Glascher, 2020). The process-specific effects of Losartan on distinct nodes may reflect that the RAS plays a complex role in regulating social reward and punishment processes.

Social deficits such as decreased social motivation or a hypersensitivity to social punishment represent a core symptom across several mental disoders including depression (Russo and Nestler, 2013; Zhang et al., 2020), SAD (Cremers et al., 2014), PTSD (Nawijn et al., 2017; Fenster et al., 2018), ASD (Delmonte et al., 2013), and schizophrenia (Mow et al., 2020). Together with accumulating evidence from previous studies (Reinecke et al., 2018; Pulcu et al., 2019; Zhou et al., 2019a) our findings suggest that Losartan may have a promising potential to enhance social motivation to obtain rewards while decreasing sensitivity to punishment in social contexts and attenuate these dysregulations in patient populations. However, long-term effects Losartan on the ambivalence toward social punishment need to be cautiously investigated.

Although the current study employed a strict pre-registered and randomized-controlled design the findings and interpretation need to be considered with the following limitations. First, research in humans has only recently begun to explore the role of the renin-angiotensin system in cognitive, emotional and reward-related processes and the underlying neurobiological mechanisms (Reinecke et al., 2018; Zhou et al., 2019a). An overarching framework of the regulatory role of the renin-angiotensin system in these domains is therefore currently lacking which limits the mechanistic interpretation of the present findings. Second, due to the proof-of-concept design the study was conducted in healthy individuals. Although this allowed us to control for a range of potential confounders, effects in patients and on the symptomatic level need to be systematically examined (Reinecke et al., 2018). Third, although the findings suggest that interactions between the RAS and the DA system may have contributed to the observed effect no direct measures of DA functioning were assessed and future molecular imaging studies are need. Chronic and dose-dependent effects of Losartan on midbrain-striatal-frontal circuits, as well as alternative mechanisms, including a contribution of systemic effects of losartan on lipid and glucose metabolism (Schupp et al., 2006), remain to be addressed in future studies.

Fourth, an additional analysis found that the guess accuracy for the correct treatment was significantly higher in the placebo group (see supplement), which indicates that most of participants were considering to be under placebo. Although an additional analysis did not reveal interaction effects between treatment guess and treatment the high rates of placebo treatment expectation in both groups may have led to an attenuation of the treatment effects. Finally, the lack of a non-social condition limits the interpretation in terms of social reward-specific effects. Future studies should investigate the interaction between drug (Losartan vs. placebo) and stimulus type (social vs. non-social). With respect to social feedback future studies may explore treatment effects on different social stimuli such as emotional faces as well as unexpected changes in people’s attributes and behavior.

In conclusion the present findings demonstrate that targeting the RAS via Losartan modulates the VTA-striatal-frontal cicruits during social feedback processing. Losartan modulated the motivational significance of social reward vs punishment feedback and concomitantly modulated the VS-prefrontal pathways. During the outcome phase Losartan attenuated VTA-insula coupling during social punishment suggesting attenuated sensitivity to social punishment. Together with the excellent safety profile of Losartan the findings may suggest a therapeutic property to enhance social motivation and attenuate the impact of negative social feedback.

## Acknowledgments and author contributions

This study was supported by the National Key Research and Development Program of China (2018YFA0701400). Data availiability: unthresholded group-level statistical maps are available via the OSF (https://osf.io/mnda4/) other data of this study is available from the corresponding author upon reasonable request. The authors report no biomedical financial interests or potential conflicts of interest.

XZ and BB conceptualized and designed the experiment. TX, YZ, RZ, and ZQ acquired data. XZ analyzed the data. XZ and BB interpreted the data and drafted the paper. XZ, BB, WZ and KK revised the manuscript critically for intellectual content. All authors commented on and gave final approval to the final version of the paper.

## References

Averbeck BB, Costa VD (2017) Motivational neural circuits underlying reinforcement learning. Nat Neurosci 20:505–512.

Barratt ES (1959) Anxiety and impulsiveness related to psychomotor efficiency. perceptual and motor skills.

Bernstein DP, Stein JA, Newcomb MD, Walker E, Pogge D, Ahluvalia T, Stokes J, Handelsman L, Medrano M, Desmond D, Zule W (2003) Development and validation of a brief screening version of the Childhood Trauma Questionnaire. Child Abuse Negl 27:169–190.

Birn RM, Roeber BJ, Pollak SD (2017) Early childhood stress exposure, reward pathways, and adult decision making. Proc Natl Acad Sci U S A 114:13549–13554.

Brown DC, Steward LJ, Ge J, Barnes NM (1996) Ability of angiotensin II to modulate striatal dopamine release via the AT1 receptor in vitro and in vivo. Br J Pharmacol 118:414–420.

Chai SY, Bastias MA, Clune EF, Matsacos DJ, Mustafa T, Lee JH, McDowall SG, Mendelsohn FAO, Albiston AL, Paxinos G (2000) Distribution of angiotensin IV binding sites (AT4 receptor) in the human forebrain, midbrain and pons as visualised by in vitro receptor autoradiography. Journal of Chemical Neuroanatomy 20:339–348.

Cremers HR, Veer IM, Spinhoven P, Rombouts SA, Roelofs K (2014) Neural sensitivity to social reward and punishment anticipation in social anxiety disorder. Front Behav Neurosci 8:439.

Culman J, von Heyer C, Piepenburg B, Rascher W, Unger T (1999) Effects of systemic treatment with irbesartan and losartan on central responses to angiotensin II in conscious, normotensive rats. Eur J Pharmacol 367:255–265.

Cuthbert BN, Insel TR (2013) Toward the future of psychiatric diagnosis: the seven pillars of RDoC. BMC Med 11:126.

Delmonte S, Gallagher L, O’Hanlon E, McGrath J, Balsters JH (2013) Functional and structural connectivity of frontostriatal circuitry in Autism Spectrum Disorder. Front Hum Neurosci 7:430.

Der-Avakian A, Markou A (2012) The neurobiology of anhedonia and other reward-related deficits. Trends Neurosci 35:68–77.

Dolen G, Darvishzadeh A, Huang KW, Malenka RC (2013) Social reward requires coordinated activity of nucleus accumbens oxytocin and serotonin. Nature 501:179–184.

Esser R, Korn CW, Ganzer F, Haaker J (2021) L-DOPA modulates activity in the vmPFC, nucleus accumbens, and VTA during threat extinction learning in humans. Elife 10.

Esteban O, Markiewicz CJ, Blair RW, Moodie CA, Isik AI, Erramuzpe A, Kent JD, Goncalves M, DuPre E, Snyder M, Oya H, Ghosh SS, Wright J, Durnez J, Poldrack RA, Gorgolewski KJ (2019) fMRIPrep: a robust preprocessing pipeline for functional MRI. Nat Methods 16:111–116.

Faulkner ML, Momenan R, Leggio L (2021) A neuroimaging investigation into the role of peripheral metabolic biomarkers in the anticipation of reward in alcohol use. Drug Alcohol Depend 221:108638.

Fenster RJ, Lebois LAM, Ressler KJ, Suh J (2018) Brain circuit dysfunction in post-traumatic stress disorder: from mouse to man. Nat Rev Neurosci 19:535–551.

Friston KJ, Jezzard P, Turner R (1994) Analysis of Functional MRI Time-Series Human Brain Mapping.

Gordon EM, Laumann TO, Marek S, Newbold DJ, Hampton JM, Seider NA, Montez DF, Nielsen AM, Van AN, Zheng A, Miller R, Siegel JS, Kay BP, Snyder AZ, Greene DJ, Schlaggar BL, Petersen SE, Nelson SM, Dosenbach NUF (2021) Human Fronto-Striatal Connectivity is Organized into Discrete Functional Subnetworks.

Gordon I, Jack A, Pretzsch CM, Vander Wyk B, Leckman JF, Feldman R, Pelphrey KA (2016) Intranasal Oxytocin Enhances Connectivity in the Neural Circuitry Supporting Social Motivation and Social Perception in Children with Autism. Sci Rep 6:35054.

Greene RK, Spanos M, Alderman C, Walsh E, Bizzell J, Mosner MG, Kinard JL, Stuber GD, Chandrasekhar T, Politte LC, Sikich L, Dichter GS (2018) The effects of intranasal oxytocin on reward circuitry responses in children with autism spectrum disorder. J Neurodev Disord 10:12.

Grimm C, Balsters JH, Zerbi V (2021a) Shedding Light on Social Reward Circuitry: (Un)common Blueprints in Humans and Rodents. Neuroscientist 27:159–183.

Grimm O, Nagele M, Kupper-Tetzel L, de Greck M, Plichta M, Reif A (2021b) No effect of a dopaminergic modulation fMRI task by amisulpride and L-DOPA on reward anticipation in healthy volunteers. Psychopharmacology (Berl) 238:1333–1342.

Gu R, Huang W, Camilleri J, Xu P, Wei P, Eickhoff SB, Feng C (2019) Love is analogous to money in human brain: Coordinate-based and functional connectivity meta-analyses of social and monetary reward anticipation. Neurosci Biobehav Rev 100:108–128.

Hetu S, Luo Y, D’Ardenne K, Lohrenz T, Montague PR (2017) Human substantia nigra and ventral tegmental area involvement in computing social error signals during the ultimatum game. Soc Cogn Affect Neurosci 12:1972–1982.

Hosseini M, Alaei HA, Havakhah S, Neemati Karimooy HA, Gholamnezhad Z (2009) Effects of microinjection of angiotensin II and captopril to VTA on morphine self-administration in rats. Acta Biol Hung 60:241–252.

Hung LW, Neuner S, Polepalli JS, Beier KT, Wright M, Walsh JJ, Lewis EM, Luo L, Deisseroth K, Dolen G, Malenka RC (2017) Gating of social reward by oxytocin in the ventral tegmental area. Science 357:1406–1411.

Izuma K, Saito DN, Sadato N (2008) Processing of social and monetary rewards in the human striatum. Neuron 58:284–294.

Klugah-Brown B, Di X, Zweerings J, Mathiak K, Becker B, Biswal B (2020) Common and separable neural alterations in substance use disorders: A coordinate-based meta-analyses of functional neuroimaging studies in humans. Hum Brain Mapp 41:4459–4477.

Lawn W, Hill J, Hindocha C, Yim J, Yamamori Y, Jones G, Walker H, Green SF, Wall MB, Howes OD, Curran HV, Freeman TP, Bloomfield MA (2020) The acute effects of cannabidiol on the neural correlates of reward anticipation and feedback in healthy volunteers. J Psychopharmacol 34:969–980.

Li D, Scott L, Crambert S, Zelenin S, Eklof AC, Di Ciano L, Ibarra F, Aperia A (2012) Binding of losartan to angiotensin AT1 receptors increases dopamine D1 receptor activation. J Am Soc Nephrol 23:421–428.

Li ZH, Bains JS, Ferguson AV (1993) Functional Evidence That the Angiotensin Antagonist Losartan Crosses the Blood-Brain-Barrier in the Rat. Brain Research Bulletin 30:33–39.

Lo MW, Goldberg MR, McCrea JB, Lu H, Furtek CI, Bjornsson TD (1995) Pharmacokinetics of losartan, an angiotensin II receptor antagonist, and its active metabolite EXP3174 in humans. Clin Pharmacol Ther 58:641–649.

Luijten M, Schellekens AF, Kuhn S, Machielse MW, Sescousse G (2017) Disruption of Reward Processing in Addiction : An Image-Based Meta-analysis of Functional Magnetic Resonance Imaging Studies. JAMA Psychiatry 74:387–398.

Martins D, Rademacher L, Gabay AS, Taylor R, Richey JA, Smith DV, Goerlich KS, Nawijn L, Cremers HR, Wilson R, Bhattacharyya S, Paloyelis Y (2021) Mapping social reward and punishment processing in the human brain: A voxel-based meta-analysis of neuroimaging findings using the social incentive delay task. Neurosci Biobehav Rev 122:1–17.

Marvar PJ, Goodman J, Fuchs S, Choi DC, Banerjee S, Ressler KJ (2014) Angiotensin type 1 receptor inhibition enhances the extinction of fear memory. Biol Psychiatry 75:864–872.

Maul B, Krause W, Pankow K, Becker M, Gembardt F, Alenina N, Walther T, Bader M, Siems WE (2005) Central angiotensin II controls alcohol consumption via its AT1 receptor. FASEB J 19:1474–1481.

McLaren DG, Ries ML, Xu G, Johnson SC (2012) A generalized form of context-dependent psychophysiological interactions (gPPI): a comparison to standard approaches. Neuroimage 61:1277–1286.

Mechaeil R, Gard P, Jackson A, Rusted J (2011) Cognitive enhancement following acute losartan in normotensive young adults. Psychopharmacology (Berl) 217:51–60.

Medelsohn FAO, Jenkins TA, Berkovic SF (1993) Effects of angiotensin II on dopamine and serotonin turnover in the striatum of conscious rats. Brain Research 613:221–229.

Modi ME, Sahin M (2019) A unified circuit for social behavior. Neurobiol Learn Mem 165:106920.

Mow JL, Gandhi A, Fulford D (2020) Imaging the “social brain” in schizophrenia: A systematic review of neuroimaging studies of social reward and punishment. Neurosci Biobehav Rev 118:704–722.

Murugan M, Jang HJ, Park M, Miller EM, Cox J, Taliaferro JP, Parker NF, Bhave V, Hur H, Liang Y, Nectow AR, Pillow JW, Witten IB (2017) Combined Social and Spatial Coding in a Descending Projection from the Prefrontal Cortex. Cell 171:1663–1677 e1616.

Narayanaswami V, Somkuwar SS, Horton DB, Cassis LA, Dwoskin LP (2013) Angiotensin AT1 and AT2 receptor antagonists modulate nicotine-evoked [(3)H]dopamine and [(3)H]norepinephrine release. Biochem Pharmacol 86:656–665.

Nawijn L, van Zuiden M, Koch SB, Frijling JL, Veltman DJ, Olff M (2017) Intranasal oxytocin increases neural responses to social reward in post-traumatic stress disorder. Soc Cogn Affect Neurosci 12:212–223.

Ohtawa M, Takayama F, Saitoh K, Yoshinaga T, Nakashima M (1993) Pharmacokinetics and biochemical efficacy after single and multiple oral administration of losartan, an orally active nonpeptide angiotensin II receptor antagonist, in humans. Br J Clin Pharmacol 35:290–297.

Pessiglione M, Seymour B, Flandin G, Dolan RJ, Frith CD (2006) Dopamine-dependent prediction errors underpin reward-seeking behaviour in humans. Nature 442:1042–1045.

Pulcu E, Shkreli L, Holst CG, Woud ML, Craske MG, Browning M, Reinecke A (2019) The Effects of the Angiotensin II Receptor Antagonist Losartan on Appetitive Versus Aversive Learning: A Randomized Controlled Trial. Biol Psychiatry 86:397–404.

Quintana DS, Lischke A, Grace S, Scheele D, Ma Y, Becker B (2021) Advances in the field of intranasal oxytocin research: lessons learned and future directions for clinical research. Mol Psychiatry 26:80–91.

Rademacher L, Krach S, Kohls G, Irmak A, Grunder G, Spreckelmeyer KN (2010) Dissociation of neural networks for anticipation and consumption of monetary and social rewards. Neuroimage 49:3276–3285.

Reinecke A, Browning M, Klein Breteler J, Kappelmann N, Ressler KJ, Harmer CJ, Craske MG (2018) Angiotensin Regulation of Amygdala Response to Threat in High-Trait-Anxiety Individuals. Biol Psychiatry Cogn Neurosci Neuroimaging 3:826–835.

Russo SJ, Nestler EJ (2013) The brain reward circuitry in mood disorders. Nat Rev Neurosci 14:609–625.

Schupp M, Lee LD, Frost N, Umbreen S, Schmidt B, Unger T, Kintscher U (2006) Regulation of peroxisome proliferator-activated receptor gamma activity by losartan metabolites. Hypertension 47:586–589.

Sharpe MJ, Chang CY, Liu MA, Batchelor HM, Mueller LE, Jones JL, Niv Y, Schoenbaum G (2017) Dopamine transients are sufficient and necessary for acquisition of model-based associations. Nat Neurosci 20:735–742.

Shkreli L, Woud ML, Ramsbottom R, Rupietta AE, Waldhauser GT, Kumsta R, Reinecke A (2020) Angiotensin involvement in trauma processing-exploring candidate neurocognitive mechanisms of preventing post-traumatic stress symptoms. Neuropsychopharmacology 45:507–514.

Sica DA, Gehr TW, Ghosh S (2005) Clinical pharmacokinetics of losartan. Clin Pharmacokinet 44:797–814.

Slotnick SD (2017) Cluster success: fMRI inferences for spatial extent have acceptable false-positive rates. Cogn Neurosci 8:150–155.

Spielberger C, Goruch R, Lushene R, Vagg P, Jacobs G (1983) Manual for the state-trait inventory STAI (form Y). Mind Garden, Palo Alto, CA, USA.

Suzuki S, Lawlor VM, Cooper JA, Arulpragasam AR, Treadway MT (2021) Distinct regions of the striatum underlying effort, movement initiation and effort discounting. Nat Hum Behav 5:378–388.

Swiercz AP, Iyer L, Yu Z, Edwards A, Prashant NM, Nguyen BN, Horvath A, Marvar PJ (2020) Evaluation of an angiotensin Type 1 receptor blocker on the reconsolidation of fear memory. Transl Psychiatry 10:363.

Torrubia R, Åvila C, Moltó J, Caseras X (2001) The Sensitivity to Punishment and Sensitivity to Reward Questionnaire (SPSRQ) as a measure of Gray’s anxiety and impulsivity dimensions. Personality and Individual Differences 31:837–862.

Trutti AC, Fontanesi L, Mulder MJ, Bazin PL, Hommel B, Forstmann BU (2021) A probabilistic atlas of the human ventral tegmental area (VTA) based on 7 Tesla MRI data. Brain Struct Funct 226:1155–1167.

Uddin LQ (2015) Salience processing and insular cortical function and dysfunction. Nat Rev Neurosci 16:55–61.

Wake SJ, Izuma K (2017) A common neural code for social and monetary rewards in the human striatum. Soc Cogn Affect Neurosci 12:1558–1564.

Watson D, Clark LA, Tellegen A (1988) Development and validation of brief measures of positive and negative affect: the PANAS scales. J Pers Soc Psychol 54:1063–1070.

Zhang D, Shen J, Bi R, Zhang Y, Zhou F, Feng C, Gu R (2020) Differentiating the abnormalities of social and monetary reward processing associated with depressive symptoms. Psychol Med:1–15.

Zhang L, Glascher J (2020) A brain network supporting social influences in human decision-making. Sci Adv 6:eabb4159.

Zhou F, Li J, Zhao W, Xu L, Zheng X, Fu M, Yao S, Kendrick KM, Wager TD, Becker B (2020) Empathic pain evoked by sensory and emotional-communicative cues share common and process-specific neural representations. Elife 9.

Zhou F, Geng Y, Xin F, Li J, Feng P, Liu C, Zhao W, Feng T, Guastella AJ, Ebstein RP, Kendrick KM, Becker B (2019a) Human Extinction Learning Is Accelerated by an Angiotensin Antagonist via Ventromedial Prefrontal Cortex and Its Connections With Basolateral Amygdala. Biol Psychiatry 86:910–920.

Zhou X, Zimmermann K, Xin F, Zhao W, Derckx RT, Sassmannshausen A, Scheele D, Hurlemann R, Weber B, Kendrick KM, Becker B (2019b) Cue Reactivity in the Ventral Striatum Characterizes Heavy Cannabis Use, Whereas Reactivity in the Dorsal Striatum Mediates Dependent Use. Biol Psychiatry Cogn Neurosci Neuroimaging 4:751–762.

