## Supplementary information for "The angiotensin antagonist Losartan modulates social reward motivation and punishment sensitivity via modulating midbrain-striato-frontal circuits"

**Supplementary methods**

**Participants – exclusion criteria**

Exclusion criteria included color blindness; systolic/diastolic blood pressure >130/90 mmHg or <90/60 mmHg; current or regular substance or medication use; current or history of medical or psychiatric disorders; any endocrinological abnormalities or contraindications for LT administration and MRI. Participants were asked to abstain from caffeinated drinks on the day of the assessment (e.g., coffee, tea, energy beverages). Two participants were excluded because their baseline blood pressure was outside our predefined criteria, one participant was excluded due to technical failure during MRI acquisition.

**Social incentive delay task**

We employed a validated social incentive delay (SID) task-fMRI paradigm with a demonstrated sensitivity for pharmacological manipulations (adopted from Nawijn, van Zuiden (1)). Briefly, the paradigm presents condition-specific cues (positive, negative, neutral) which signal that a social reward can be obtained, or a social punishment can be avoided (anticipation). Next, participants undergo a reaction time task which is followed by the presentation of a performance-dependent social reward, punishment, or neutral feedback (outcome) (Figure 1B). Participants received task instructions and a practice session prior to the formal experiment. During the practice session all participants were informed about the types of outcomes that might be presented and that the corresponding outcome might or might not occur (no information on the probability was provided). During the formal paradigm 27 trials for reward and punishment conditions, and 18 trials for the neutral condition were presented (pseudo-randomized). Each trial started with presentation of a geometric cue indicating the trial type (circle: reward, triangle: punishment, and square: neutral, Figure 1C). After a delay, the target was presented in the center of screen and participants were required to press a button as fast as possible. Responses within target presentation time represented hits while omissions or responses outside of target presentation time represented misses. To facilitate a sufficient number of trials for each outcome condition an adaptive performance algorithm was employed. To this end, trial-wise reaction times (RT) were recorded in real-time and employed to adjust the duration of the next target presentation to the individual performance. To this end individual RTs were used to tailor the duration of target presentation to individual performance. The total duration consisted of the baseline time (500ms) and a change time based on the adaptive algorithm that was initially evaluated in an independent sample with comparable demographic characteristic. If the response time of the participant exceeded the target time on a given trial for the reward condition the duration of the next target in the reward condition was be increased by 60ms, in case the participant responded in time the duration of the next trial was decreased by 40ms. For the punishment condition an increase of 20ms or a decrease of 70ms was employed. The individual adaptation in terms of increasing or decreasing the target duration times allowed to yield approx. 66.7% of reward-cue and punishment-cue trials being followed by social reward or punishment, respectively. This resulted in a performance-dependent sufficient number of reward (hit) feedback trials (±18) and punishment (miss) feedback trials (±18) for further analyses. The target was followed by an adaptive inter-trial-interval and presentation of the condition- and performance-dependent outcome. In the social reward condition hits resulted in rewarding social feedback, i.e. a smiling person in thumbs-up pose, while misses resulted in neutral feedback, i.e. scrambled picture of the person. In the social punishment condition hits allowed avoidance of social punishment (neutral feedback), while misses resulted in social punishment feedback, i.e. a person with a contemptuous look in thumbs-down pose. For neutral trials both, hits and misses resulted in neutral feedback. The experimental materials for the outcome were initially collected and rated by an independent sample. To further explore the subsequent emotional effects of the paradigm participants rated perceived arousal, likeability, dislikeability, intensity, valence of cues and outcomes on a 9-point Likert scale after the fMRI session (Figure 1A).

**Behavioral analysis**

To maintain the full information of the trial structure in the SID paradigm and to increase sensitivity of the analysis a linear mixed model (R package ‘lme4’) was used with condition (social reward, social punishment, and neutral) and treatment (losartan and placebo) as two fixed factors and with subject as a random factor to account for individual changes in reaction time (RT).

The ANOVA on the affective ratings of the cues was conducted employing the R package ‘afex’ (https://CRAN.R-project.org/package=afex) with condition (reward, punishment, and neutral) and time (before and after experiment) as within-subject factors, and treatment (Losartan and placebo) as between-subjects factor. To match the RT analysis and enhance the sensitivity a liner mixed model was used to analyze the ratings of outcome cues with condition (social reward, social punishment, and neutral control) and treatment (Losartan and placebo) as fixed factors and subject as a random factor.

**MRI data acquisition**

MRI data was acquired using a 3.0 Tesla GE MR750 system (General Electric Medical System, Milwaukee, WI, USA). T1-weight high-resolution anatomical images were acquired with a spoiled gradient echo pulse sequence, repetition time (TR) = 6 ms, echo time (TE) = 2 ms, flip angle = 12°, field of view (FOV) = 256 × 256 mm, acquisition matrix = 256 × 256, slice thickness = 1 mm, voxel size = 1 × 1 × 1 mm. Functional data was acquired using a T2*-weight Echo Planar Imaging (EPI) sequence with the following parameters: TR = 2000 ms, TE = 30 ms, flip angle = 90°, acquisition matrix = 64 × 64, slice thickness = 3 mm, and 39 slices with an interleaved ascending order. Head movements were minimized by using comfortable head cushions. For each participant 318 volumes were collected.

**MRI data preprocessing**

MRI data processing in this manuscript are based on fMRIPrep 20.2.1 (RRID:SCR_016216) (2), which is based on Nipype 1.5.1 (RRID:SCR_002502) (3).

**Anatomical data preprocessing**

A total of T1-weighted (T1w) images were found within the input BIDS dataset. The T1-weighted (T1w) image was corrected for intensity non-uniformity (INU) with N4BiasFieldCorrection (4), distributed with ANTs 2.3.3 (RRID:SCR_004757), and used as T1w-reference throughout the workflow. The T1w-reference was then skull-stripped with a Nipype implementation of the antsBrainExtraction.sh workflow (from ANTs), using OASIS30ANTs as target template. Brain tissue segmentation of cerebrospinal fluid (CSF), white-matter (WM) and gray-matter (GM) was performed on the brain-extracted T1w using fast (FSL 5.0.9, RRID:SCR_002823) (5). Brain surfaces were reconstructed using recon-all (FreeSurfer 6.0.1, RRID:SCR_001847), and the brain mask estimated previously was refined with a custom variation of the method to reconcile ANTs-derived and FreeSurfer-derived segmentations of the cortical gray-matter of Mindboggle (RRID:SCR_002438) (6). Volume-based spatial normalization to the standard space (MNI152NLin6Asym) was performed through nonlinear registration with antsRegistration (ANTs 2.3.3), using brain-extracted versions of both T1w reference and the T1w template. The following templates were selected for spatial normalization: FSL’s MNI ICBM 152 non-linear 6th Generation Asymmetric Average Brain Stereotaxic Registration Model (RRID:SCR_002823; TemplateFlow ID: MNI152NLin6Asym) (7).

**Functional data preprocessing**

For each of the two BOLD runs found per subject (across all tasks and sessions), the following preprocessing was performed. First, a reference volume and its skull-stripped version were generated using a custom methodology of fMRIPrep. Susceptibility distortion correction (SDC) was omitted. The BOLD reference was then co-registered to the T1w reference using bbregister (FreeSurfer) which implements boundary-based registration (8). Co-registration was configured with six degrees of freedom. Head-motion parameters with respect to the BOLD reference (transformation matrices, and six corresponding rotation and translation parameters) are estimated before any spatiotemporal filtering using mcflirt (FSL 5.0.9) (9). BOLD runs were slice-time corrected using 3dTshift from AFNI 20160207 (RRID:SCR_005927) (10). The BOLD time-series (including slice-timing correction when applied) were resampled onto their original, native space by applying the transforms to correct for head-motion. These resampled BOLD time-series will be referred to as preprocessed BOLD in original space, or just preprocessed BOLD. The BOLD time-series were resampled into several standard spaces, correspondingly generating the following spatially-normalized, preprocessed BOLD runs: MNI152NLin6Asym. First, a reference volume and its skull-stripped version were generated using a custom methodology of fMRIPrep. The head-motion estimates calculated in the correction step were placed within the corresponding confounds file. The confound time series derived from head motion estimates and global signals were expanded with the inclusion of temporal derivatives and quadratic terms for each (11). All resamplings can be performed with a single interpolation step by composing all the pertinent transformations (i.e. head-motion transform matrices, susceptibility distortion correction when available, and co-registrations to anatomical and output spaces). Gridded (volumetric) resamplings were performed using antsApplyTransforms (ANTs), configured with Lanczos interpolation to minimize the smoothing effects of other kernels (12). Non-gridded (surface) resamplings were performed using mri_vol2surf (FreeSurfer).

Many internal operations of fMRIPrep use Nilearn 0.6.2 (RRID:SCR_001362) (13), mostly within the functional processing workflow. For more details of the pipeline, see the section corresponding to workflows in fMRIPrep’s documentation.

**Individual-level fMRI model**

On the first-level the SID task was modeled employing the hemodynamic response function (HRF) and corresponding derivatives on the onsets of the experimental conditions, i.e. cue, target, and outcome. The three different types of anticipation were modeled according to the cues signaling a potential social reward, social punishment, and neutral outcome. In line with previous studies the cue and subsequent delay were modelled as anticipation period (14–17). In line with the potential response pattern of the participants five different types of outcome were modeled, including reward or neutral feedback in reward trial, punishment or neutral feedback in punishment trial, and neutral feedback in neutral trial. Additional confound regressors in the SPM statistical model included target responses and head motion as defined by six rigid movement parameters from the motion correction.

**Exploratory Functional Connectivity Analysis**

Based on our a priori regional hypotheses the VTA, VS and DS masks were employed as seed regions for the analyses of task-modulated functional connectivity. To this end generalized context-dependent psychophysiological interaction analyses (gPPI) were computed as implemented in the gPPI toolbox (<http://www.nitrc.org/projects/gppi>, RRID: SCR_009489) (18). Combined with the a priori defined VS/DS/VTA masks and neural activity during receipt of feedback, three peak coordinates (VS: [22/-6/-10], DS: [-14/-2/-8], VTA: [10/-14/-12]) were employed as seed regions and the extracted BOLD time series from the subregion-specific masks were treated as main physiological factors in the gPPI first level design matrix. In line with the BOLD level analysis session-specific regressors for the experimental conditions (reward, punishment) and the other controlled regressors same as activation analysis were additionally included in the first level design matrix and motion parameters were included to improve motion control.

**Supplementary results**

**Affective impact of the experimental manipulation**

With respect to cue ratings, the ANOVA revealed significant main effects of arousal (time, *F* = 17.04, *p* < 0.001; condition, *F* = 13.58, *p* < 0.001, Table S1), charm (time, *F* = 17.04, *p* < 0.001; condition, *F* = 13.58, *p* < 0.001), dislikeability (condition, *F* = 15.98, *p* < 0.001), intensity (time, *F* = 35.85, *p* < 0.001; condition, *F* = 10.42, *p* < 0.001), valence (condition, *F* = 18.27, *p* < 0.001), and interaction effects of condition and time on charm (*F* = 6.37, *p* = 0.002) and dislikeability (*F* = 3.22, *p* = 0.044). The significant effect of time indicated that the experiment successfully induced stimulus-outcome associations. Regarding to outcome ratings, the linear mixed model revealed significant main effects of condition on charm (*F* = 689.416, *p* < 0.0001), intensity (*F* = 357.869, *p* < 0.0001), and valence (*F* = 812.378, *p* < 0.0001).

**Neural activity during the anticipation phase**

The ANOVA analyses were conducted on the extracted parameter estimated from a priori ROIs including VTA (treatment: *F* = 2.67, *p* = 0.106, condition: *F* = 0.75, *p* = 0.467, interaction: *F* = 0.31, *p* = 0.724), VS (treatment: *F* = 1.28, *p* = 0.262, condition: *F* = 5.40, *p* = 0.007, interaction: *F* = 0.22, *p* = 0.780), and DS (treatment: *F* = 0.96, *p* = 0.330, condition: *F* = 2.60, *p* = 0.080, interaction: *F* = 0.65, *p* = 0.516). Only significant main effect of condition was observed in VS, and the post hoc test showed lower response to reward relative to the neutral condition (*t* = 3.257, *p* = 0.0039).

In addition, an exploratory whole-brain analysis confirmed the absence of significant treatment main and interaction effects. Only significant condition effects were observed at whole-brain cluster level *p_FWE_* < 0.05 (initial formatting uncorrected *p* < 0.001) during anticipation phase (Figure S1A and Table S2).

**Neural activity during outcome phase**

The ANOVA analyses were conducted on the extracted parameter estimated from a priori ROIs including VTA (treatment: *F* = 0.03, *p* = 0.872, condition: *F* = 0.73, *p* = 0.477, interaction: *F* = 3.24, *p* = 0.043), VS (treatment: *F* = 0.53, *p* = 0.468, condition: *F* = 40.50, *p* < 0.001, interaction: *F* = 0.51, *p* = 0.599), and DS (treatment: *F* = 1.67, *p* = 0.199, condition: *F* = 9.10, *p* < 0.001, interaction: *F* = 0.19, *p* = 0.822). For both VS and DS, the post hoc tests showed higher responses to reward relative to the neural condition (VS, *t* = 7.158, *p* < 0.0001; DS, *t* = 2.985, *p* = 0.0091) and punishment condition (VS, *t* = 8.304, *p* < 0.0001; DS, *t* = 4.133, *p* = 0.0002).

In addition, exploratory whole-brain analysis confirmed the absence of significant treatment main and interaction effects. Only significant condition effects were observed at whole-brain cluster level *p_FWE_* < 0.05 (initial formatting uncorrected *p* < 0.001) during the outcome phase (Figure S1B and Table S2).

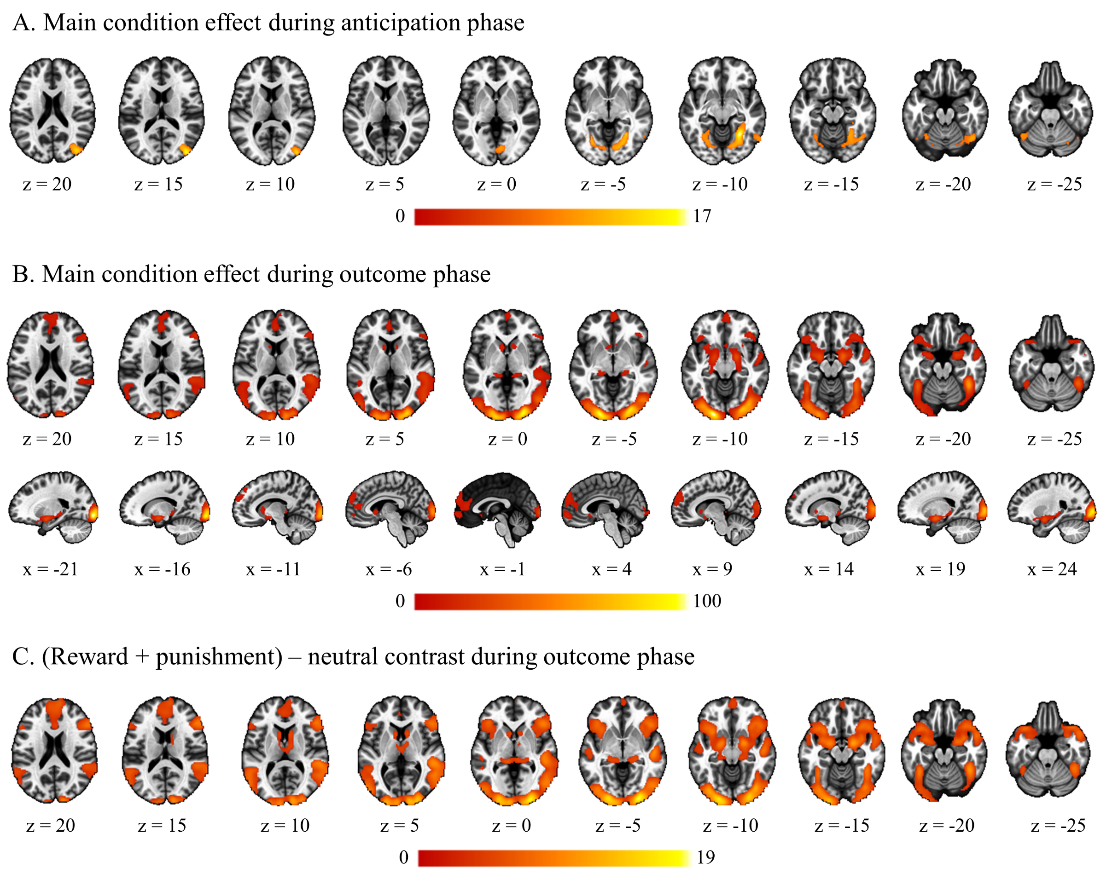

Figure S1. Brain activity during SID task. All thresholds applied on the whole-brain cluster level *p_FWE_* < 0.05 (initial cluster forming threshold uncorrected p < 0.001).

Table S1. Losartan effects on reaction time and affective response to outcome stimuli

|  | Sum Sq | Mean Sq | NumDF | DenDF | F | p |
| --- | --- | --- | --- | --- | --- | --- |
| Reaction time |  |  |  |  |  |  |
| Treatment | 5836.184 | 5836.184 | 1 | 85.0928 | 1.097 | 0.298 |
| Condition | 27729.964 | 13864.982 | 2 | 5983.9603 | 2.607 | 0.074 |
| Interaction | 39417.958 | 19708.979 | 2 | 5983.9603 | 3.706 | 0.025 |
| Arousal |  |  |  |  |  |  |
| Treatment | 1.166 | 1.166 | 1 | 85 | 0.466 | 0.497 |
| Condition | 2028.444 | 1014.222 | 2 | 1997 | 404.983 | <.0001 |
| Interaction | 24.614 | 12.307 | 2 | 1997 | 4.914 | 0.007 |
| Charm |  |  |  |  |  |  |
| Treatment | 0.014 | 0.014 | 1 | 85 | 0.005 | 0.943 |
| Condition | 3697.834 | 1848.917 | 2 | 1997 | 689.416 | <.0001 |
| Interaction | 7.447 | 3.723 | 2 | 1997 | 1.388 | 0.250 |
| Dislikeability |  |  |  |  |  |  |
| Treatment | 0.285 | 0.285 | 1 | 85 | 0.087 | 0.769 |
| Condition | 4166.394 | 2083.197 | 2 | 1997 | 633.848 | <.0001 |
| Interaction | 22.434 | 11.217 | 2 | 1997 | 3.413 | 0.033 |
| Intensity |  |  |  |  |  |  |
| Treatment | 3.201 | 3.201 | 1 | 85 | 1.197 | 0.277 |
| Condition | 1913.460 | 956.730 | 2 | 1997 | 357.869 | <.001 |
| Interaction | 14.011 | 7.006 | 2 | 1997 | 2.621 | 0.073 |
| Valence |  |  |  |  |  |  |
| Treatment | 0.647 | 0.647 | 1 | 85 | 0.300 | 0.586 |
| Condition | 3506.816 | 1753.408 | 2 | 1997 | 812.378 | <.0001 |
| Interaction | 2.046 | 1.023 | 2 | 1997 | 0.474 | 0.623 |

Table S2. Brain activity during SID task.

| Cluster region | Cluster size | x | y | z | *F/t* value |
| --- | --- | --- | --- | --- | --- |
| Anticipation phase | |  |  |  |  |
| Occipital lobe, parahippocampa gyrus, fusiform, lingual gyrus | 2828 | 28 | -56 | -10 | 17.46 |
|  |  | 44 | -62 | -18 | 11.71 |
|  |  | -26 | -70 | -6 | 11.64 |
| Occipital lobe, MTG | 647 | 42 | -86 | 14 | 16.69 |
| Outcome phase | |  |  |  |  |
| Occipital lobe, temporal lobe | 13431 | -20 | -98 | -8 | 107.19 |
|  |  | 26 | -96 | 0 | 96.68 |
|  |  | 28 | -90 | -6 | 91.50 |
| IFG, MFG, STG, insula, subcortical regions* | 4928 | 20 | -2 | -14 | 39.05 |
|  |  | -18 | -4 | -14 | 31.17 |
|  |  | 54 | 32 | 12 | 18.91 |
| MFG, SFG, anterior cingulate | 2132 | 4 | 62 | -8 | 15.07 |
|  |  | -2 | 46 | 12 | 13.86 |
|  |  | 6 | 62 | 34 | 13.32 |
| (Reward + punishment) > neutral | |  |  |  |  |
| Occipital lobe, STG, MTG, IFG, MFG, subcortical regions* | 30964 | 26 | -96 | -4 | 18.53 |
|  |  | -22 | -96 | -10 | 16.52 |
|  |  | 14 | -100 | 6 | 13.87 |
| SFG, MFG, anterior cingulate, supplementary motor area | 6288 | 2 | 52 | 34 | 8.69 |
|  |  | 8 | 60 | 36 | 8.10 |
|  |  | 8 | 22 | 60 | 7.63 |
| Cerebellum | 456 | -4 | -58 | -38 | 7.06 |
| MFG | 330 | 2 | 62 | -10 | 5.08 |

Note: All clusters passed the threshold at whole-brain cluster level *p_FWE_* < .05. IFG = inferior frontal gyrus, SFG = superior frontal gyrus, MFG = middle frontal gyrus, STG = superior temporal gyrus, MTG = middle temporal gyrus. * parahippocampa gyrus, caudate, putamen, amygdala, thalamus.

Table S3. Subregional location of interaction effects on functional connectivity

| Anticipation phase, VS seed | |  |  |  |  |
| --- | --- | --- | --- | --- | --- |
| Cluster 1, k=309 | Assignment based on MPM | Percent of cluster volume in area | Percent of area activated by cluster |  |  |
|  | Area Fo5 | 4.2 | 3.9 |  |  |
|  | Area Fp1 | 1.4 | 0.2 |  |  |
|  | - | - | - |  |  |
|  | Probability exceedance by Area | Mean probability at Cluster location (%) | Mean probability across PMap (%) | Ratio | 95%CI |
|  | Area Fo5 | 16.8 | 19.5 | 0.86 | 0.80-0.91 |
|  | Area Fp1 | 9.4 | 39.5 | 0.24 | 0.22-0.26 |
|  | Area Fo6 | 0 | 26.0 | 0 | 0 |
| Outcome phase, VTA seed | |  |  |  |  |
| Cluster 1, k=427 | Assignment based on MPM | Percent of cluster volume in area | Percent of area activated by cluster |  |  |
|  | Area Id2 | 12.5 | 35.6 |  |  |
|  | Area Id5 | 9.2 | 17.3 |  |  |
|  | Area Ig2 | 7.9 | 16.9 |  |  |
|  | Probability exceedance by Area | Mean probability at Cluster location (%) | Mean probability across PMap (%) | Ratio | 95%CI |
|  | Area Id2 | 25.1 | 14.7 | 1.70 | 1.60-1.79 |
|  | Area Id5 | 30.9 | 20.2 | 1.53 | 1.44-1.62 |
|  | Area Ig2 | 20.7 | 17.3 | 1.19 | 1.11-1.26 |
| Cluster 2, k=246 | Assignment based on MPM | Percent of cluster volume in area | Percent of area activated by cluster |  |  |
|  | Area Id6 | 18.9 | 6.8 |  |  |
|  | Area Id5 | 7.1 | 7.7 |  |  |
|  | Area Ia | 2.5 | 6.1 |  |  |
|  | Probability exceedance by Area | Mean probability at Cluster location (%) | Mean probability across PMap (%) | Ratio | 95%CI |
|  | Area Id6 | 34.1 | 24.9 | 1.37 | 1.30-1.44 |
|  | Area Id5 | 23.4 | 20.2 | 1.16 | 1.08-1.23 |
|  | Area Ia | 15.4 | 19.5 | 0.79 | 0.68-0.89 |
| Cluster 3, k=213 | Assignment based on MPM | Percent of cluster volume in area | Percent of area activated by cluster |  |  |
|  | - | - | - |  |  |
|  | Probability exceedance by Area | Mean probability at Cluster location (%) | Mean probability across PMap (%) | Ratio | 95%CI |
|  | Area 6d2 | 6.2 | 19.1 | 0.32 | 0.30-0.35 |
|  | Area 6d3 | 4.1 | 20.4 | 0.20 | 0.19-0.21 |
|  | Area 6mr/preSMA | 0.3 | 24.6 | 0.01 | 0.01-0.02 |

Only the first 3 areas for each term were presented. MPM = Maximum Probability Map, lateral orbitofrontal cortex: Area Fo5, Area Fo6, frontal pole: Area Fp1, dysgranular insula: Area Id2, Area Id5, Area Id6, granular insula: Area Ig2, agranular insula: Area Ia, dorsal precentral gyrus: Area 6d2, superior frontal sulcus: Area 6d3, posterior medial superior frontal gyrus: Area 6mr/preSMA

**References**

1. Nawijn L, van Zuiden M, Koch SBJ, Frijling JL, Veltman DJ, Olff M (2017): Intranasal oxytocin increases neural responses to social reward in post-traumatic stress disorder. *Soc Cogn Affect Neurosci* 12: 212–223.

2. Esteban O, Markiewicz CJ, Blair RW, Moodie CA, Isik AI, Erramuzpe A, *et al.* (2019): fMRIPrep: a robust preprocessing pipeline for functional MRI. *Nat Methods* 16: 111–116.

3. Gorgolewski K, Burns CD, Madison C, Clark D, Halchenko YO, Waskom ML, Ghosh SS (2011): Nipype: A flexible, lightweight and extensible neuroimaging data processing framework in Python. *Front Neuroinform* 5: 13.

4. Tustison NJ, Avants BB, Cook PA, Zheng Y, Egan A, Yushkevich PA, Gee JC (2010): N4ITK: Improved N3 bias correction. *IEEE Trans Med Imaging* 29: 1310–1320.

5. Zhang Y, Brady M, Smith S (2001): Segmentation of brain MR images through a hidden Markov random field model and the expectation-maximization algorithm. *IEEE Trans Med Imaging* 20: 45–57.

6. Klein A, Ghosh SS, Bao FS, Giard J, Häme Y, Stavsky E, *et al.* (2017): Mindboggling morphometry of human brains. *PLoS Comput Biol* 13. https://doi.org/10.1371/journal.pcbi.1005350

7. Evans AC, Janke AL, Collins DL, Baillet S (2012): Brain templates and atlases. *Neuroimage* 62: 911–922.

8. Greve DN, Fischl B (2009): Accurate and robust brain image alignment using boundary-based registration. *Neuroimage* 48: 63–72.

9. Jenkinson M, Bannister P, Brady M, Smith S (2002): Improved Optimization for the Robust and Accurate Linear Registration and Motion Correction of Brain Images. *Neuroimage* 17: 825–841.

10. Cox RW, Hyde JS (1997): Software tools for analysis and visualization of fMRI data. *NMR Biomed* 10: 171–178.

11. Satterthwaite TD, Elliott MA, Gerraty RT, Ruparel K, Loughead J, Calkins ME, *et al.* (2013): An improved framework for confound regression and filtering for control of motion artifact in the preprocessing of resting-state functional connectivity data. *Neuroimage* 64: 240–256.

12. Lanczos C (1964): Evaluation of Noisy Data. *J Soc Ind Appl Math Ser B Numer Anal* 1: 76–85.

13. Abraham A, Pedregosa F, Eickenberg M, Gervais P, Mueller A, Kossaifi J, *et al.* (2014): Machine learning for neuroimaging with scikit-learn. *Front Neuroinform* 8. https://doi.org/10.3389/fninf.2014.00014

14. Zhang D, Shen J, Bi R, Zhang Y, Zhou F, Feng C, Gu R (2020): Differentiating the abnormalities of social and monetary reward processing associated with depressive symptoms. *Psychol Med*. https://doi.org/10.1017/S0033291720003967

15. Faulkner ML, Momenan R, Leggio L (2021): A neuroimaging investigation into the role of peripheral metabolic biomarkers in the anticipation of reward in alcohol use. *Drug Alcohol Depend* 221. https://doi.org/10.1016/j.drugalcdep.2021.108638

16. Lawn W, Hill J, Hindocha C, Yim J, Yamamori Y, Jones G, *et al.* (2020): The acute effects of cannabidiol on the neural correlates of reward anticipation and feedback in healthy volunteers. *J Psychopharmacol* 34: 969–980.

17. Rademacher L, Krach S, Kohls G, Irmak A, Gründer G, Spreckelmeyer KN (2010): Dissociation of neural networks for anticipation and consumption of monetary and social rewards. *Neuroimage* 49: 3276–3285.

18. McLaren DG, Ries ML, Xu G, Johnson SC (2012): A generalized form of context-dependent psychophysiological interactions (gPPI): A comparison to standard approaches. *Neuroimage* 61: 1277–1286.
